# Integrated analysis of human transcriptome data for Rett syndrome finds a network of involved genes

**DOI:** 10.1101/274258

**Authors:** Friederike Ehrhart, Susan L. Coort, Lars Eijssen, Elisa Cirillo, Eric E. Smeets, Nasim Bahram Sangani, Chris T. Evelo, Leopold M.G. Curfs

## Abstract

Rett syndrome (RTT) is a rare disorder causing severe intellectual and physical disability. The cause is a mutation in the gene coding for the methyl-CpG binding protein 2 (MECP2), a multifunctional regulator protein. Purpose of the study was integration and investigation of multiple gene expression profiles in human cells with impaired *MECP2* gene to obtain a data-driven insight in downstream effects. Information about changed gene expression was extracted from five previously published studies. We identified a set of genes which are significantly changed not in all but several transcriptomics datasets and were not mentioned in the context of RTT before. Using overrepresentation analysis of molecular pathways and gene ontology we found that these genes are involved in several processes and molecular pathways known to be affected in RTT. Integrating transcription factors we identified a possible link how MECP2 regulates cytoskeleton organization via MEF2C and CAPG. Integrative analysis of omics data and prior knowledge databases is a powerful approach to identify links between mutation and phenotype especially in rare disease research where little data is available.

**Abbreviations:** Rett syndrome (RTT), embryonic stem cells (ESCs), induced pluripotent stem cells (iPSCs), fold change (FC), Gene Ontology (GO), EIF (eukaryotic initiation of transcription factor)

For genes the symbols according to the HGNC nomenclature were used.

## Introduction

Rett syndrome (RTT, MIM #312750) is a neurological disorder which was first described in 1966 by A. Rett (Rett, 1966), an Austrian pediatrician who observed the same stereotypic hand movements in some of his patients. It is a rare disease which affects between 1:10,000 to 1:20,000 live born children and is predominantly seen in females (Laurvick et al, 2006). The symptoms are stagnation, followed by regression and loss of motor and communication skills between the age of 6 – 18 months after almost normal pre- and postnatal development (Liyanage & Rastegar, 2014). Neuronal function and development are strongly impaired. Patients develop intellectual disability, motor impairments, and characteristic hand stereotypes (e.g. hand washing). Often, but not always, abnormal breathing patterns (Viemari et al, 2005), sleep disorders (Spruyt & Curfs, 2015), epilepsy, scoliosis, growth deficit, cardiac dysrhythmias (Hara et al, 2015), constipation and dystonia are observed (Liyanage & Rastegar, 2014). In later stages the motoric abilities typically decrease while cognitive and communication skills are stable and often even improving.

RTT is linked to a mutation in the gene encoding methyl-CpG binding protein 2 (MECP2) which is located on the X-chromosome (Amir et al, 1999). MECP2 is an important regulator of neuronal function and development. It influences neuronal differentiation, maturation, morphology and synaptic plasticity and its RNA and protein are found in almost all tissues (Uhlen et al, 2015). MECP2 expression is low in fetal development but increases during postnatal development which may explain the late and gradual onset of the syndrome. Switching off *MECP2* in juvenile or adult stage in mice induced the RTT-like phenotype indicating that MECP2 is not only essential during neuronal development but also required for neurological function and maintenance (Nguyen et al, 2012). On the molecular level, MECP2 acts as regulator of gene expression and depending on the cofactor functions as activator or inhibitor. It influences alternative splicing, and miRNA expression (Chahrour et al, 2008; Ehrhart et al, 2016; Gonzales et al, 2012; Jordan et al, 2007; Liyanage & Rastegar, 2014). RTT is also a disorder which is well known for epigenetic implications because MECP2 not only binds on methylated DNA sites but is also involved in maintenance and modification of the methylation pattern of DNA (Bienvenu & Chelly, 2006; Sperlazza et al, 2017). For a previous study we created a comprehensive visualization of known MECP2 actions which can be found on WikiPathways (http://www.wikipathways.org/instance/WP3584) (Ehrhart et al, 2016). MECP2 being such a wide range influencer of gene expression, epigenetic imprinting and neurotransmitter action, makes it challenging to gather information about the downstream effects. Discovering which gene expression patterns are directly affected by an impaired *MECP2* and reconstructing the downstream pathways which finally lead to the phenotype is a still in its infancy.

Nowadays, cell biology uses high throughput methods to investigate the causes and consequences of diseases on the basis of whole metabolomes, transcriptomes and genomes, separately and integrative. The induced changes on the cellular level represent intricate biological networks including molecular interactions which finally lead to a specific disorder phenotype. By performing meta- or integrational analysis of published transcriptomics data, various studies of neurological disorders like autism-spectrum disorder (Ch’ng et al, 2015; Li et al, 2014), schizophrenia (Mistry et al, 2013), multiple sclerosis (Raddatz et al, 2014), intracranial aneurysms (Roder et al, 2012), epilepsy (Rogic & Pavlidis, 2009), and aging and age-associated spatial learning impairment (Uddin & Singh, 2013) have obtained new insights. However, to our knowledge, this approach has not been applied yet in RTT. Being able to reuse published data is a positive consequence of biological raw data being increasingly well documented, stored in data repositories, and made interoperable for reuse (Wilkinson et al, 2016).

The aim of the present study is to investigate differential gene expression and biological pathways in a collection of previously published transcriptomics studies obtained from human cells with impaired MECP2 function. The raw data was derived and integrated from five independent studies published online in GEO (Barrett et al, 2013; Edgar et al, 2002) and ArrayExpress (Kolesnikov et al, 2015) databases (Uhlen et al, 2016). From these raw datasets the differentially expressed gene lists were extracted. These were then used for secondary analysis, namely, pathway analysis (WikiPathways, using PathVisio software) (Kelder et al, 2012) and gene ontology analysis (Gene Ontology (GO) using GO-Elite software (Zambon et al, 2012)).

## Results

### Comparison of gene expression in the different samples

The differentially expressed gene lists, obtained from raw data of five previously published studies, were extracted and according to their tissue origin sorted in three different tissue groups including human brain tissue, fibroblasts, and neuronal cells (for the details see sup. Table 1 Dataset description). The three tissue groups were defined according to tissue/cell sample similarity in the different studies used: human brain (which includes the three age group samples from E-GEOD-6955), fibroblasts (E-GEOD-21037-F and all three E-MEXP-1956 samples), and neuronal cells (E-GEOD-21037-I, E-GEOD-4600, E-GEOD-50584-4W). For the brain and fibroblast samples there was only one sample per dataset available, so the log2FC was used as a threshold instead of significance (p value). There were two different log2FC thresholds investigated, ± 0.58 (fold change of ± 1.5) and a more restrictive threshold of ± 1.0 (fold change of ± 2). The numbers of differentially expressed genes per experimental group and the overlap within each of the tissue groups is shown for log2FC threshold ± 1 in figure 1 (for 0.58 see sup. Table 2: Differentially expressed gene lists, and sup. Table 3: Summary of overlapping genes, pathways and GO). The human brain group had only eleven commonly changed genes, fibroblasts as well 11, and neuronal cell culture group had 51 common differentially expressed genes. Notable is also, that the number of differentially expressed genes within the embryonic stem cell group increases from neuronal precursor (E-GEOD-50584-NP = 425) towards neuronal cells (after 2 weeks: 2W = 1313 and after 4 weeks: 4W = 2084). Fibroblast samples, in which the impaired *MECP2* and the control are collected from the same person (the E-MEXP-1952 samples), are different in 204, 269 or 303 genes (Fig 1) while the fibroblast samples, in which the impaired *MECP2* and the control are derived from different persons differ in 840 genes. Overlapping differentially expressed genes and their expression change (log2FC) are shown in sup. Table 1 and 3. Over all tissues, the genes which show differential expression in one tissue group plus in at least one more sample from another group (meaning differential expression in 5-6 experimental samples) are collected in Table 1. There were no genes which were found to be differentially expressed in more than six group comparisons for log2FC threshold 1, or more than eight group comparisons for 0.58.

**Table 1:** Genes, which are differentially expressed in 5 or 6 different experimental group comparisons (log2FC threshold for brain and fibroblast samples = ± 1).

| Gene symbol | Description | Expression direction* | Tissue expression profile according to tissue atlas | Biol. function according to gene ontology |
| --- | --- | --- | --- | --- |
| YBX3 | Y-box binding protein 3 | Brain | expressed in all, enriched in skeletal muscle | DNA binding, transcription factor |
|  |  | Fibroblasts |  |  |
|  |  | Cell culture |  |  |
| LITAF | Lipopolysaccharide induced TNF factor |  | expressed in all tissues | LPS binding and signaling, transcription factor |
| HSPA2 | Heat shock protein family A (Hsp70) member 2 |  | expressed in all tissues | HSP - protein folding |
| CDH2 | Cadherin 2 |  | adrenal gland, heart, cell adhesion, synapsis | cell adhesion, signal transduction |
| CAPG | Capping actin protein, gelsolin like |  | all tissues | capping of forked actin filaments |
| SERPING1 | Serpin family G member 1 |  | all tissues | complement cascade regulator (C1 inhibitor) |
| CCL2 | C-C motif chemokine ligand 2 |  | all tissues | immunoregulatory and inflammatory process |
| SRSF11 | Serine and arginine rich splicing factor 11 |  | all tissues | mRNA processing |
| ACAT2 | Acetyl-CoA acetyltransferase 2 |  | all tissues | lipid metabolism |
| MAP1B | Microtubule associated protein 1B |  | all tissues, enriched in cerebral cortex | microtubule assembly, neurogenesis |
| MEIS2 | Meis homeobox 2 |  | mixed | homeobox gene, cell differentiation |
| TAGLN3 | Transgelin 3 |  | brain, brain tissues | central nervous system development |
| SLC3A2 | Solute carrier family 3 member 2 |  | all tissues | amino acid transporter, Ca<sup>2+</sup> regulation |
| ABAT | 4-aminobutyrate aminotransferase |  | enriched in brain, kidney, liver | GABA metabolizer |

\* Expression direction shows whether the majority of samples in the subgroups brain, fibroblasts and cell culture are up (red) or down (blue) or do not change (white).

**Table 2:**
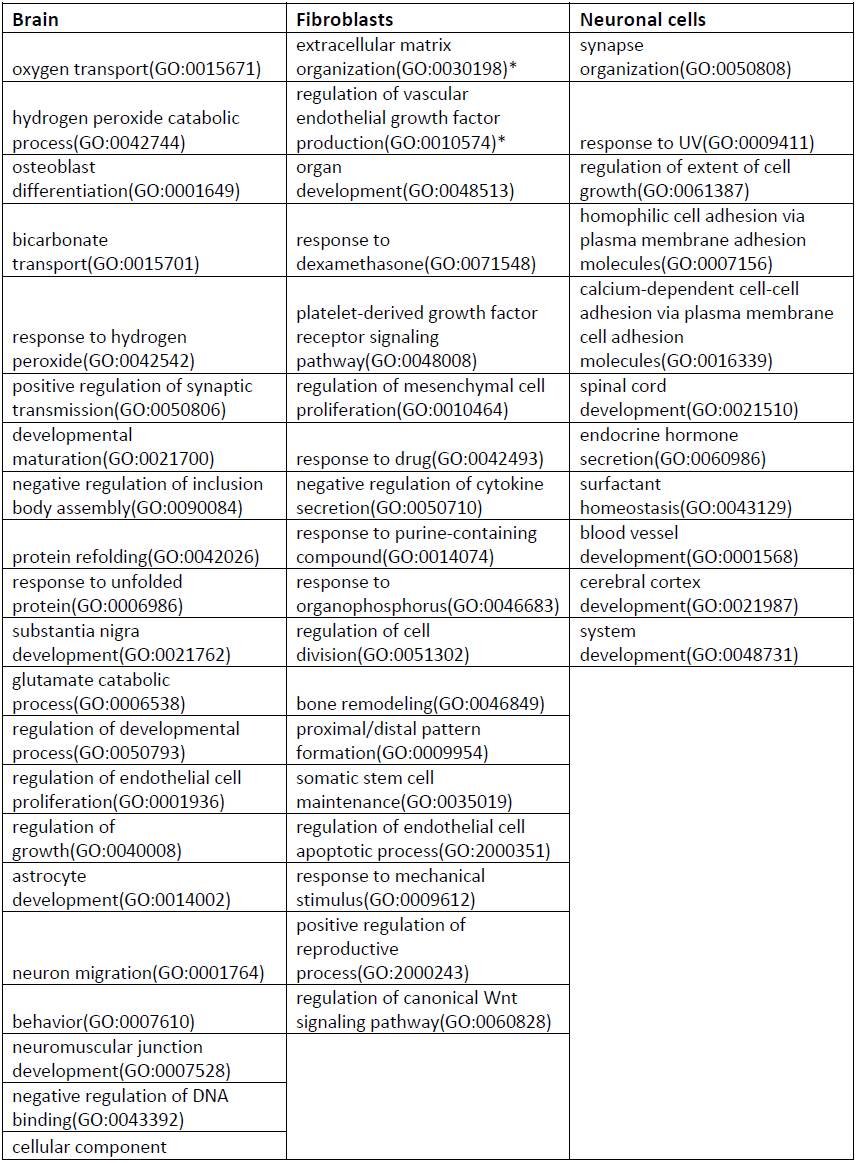
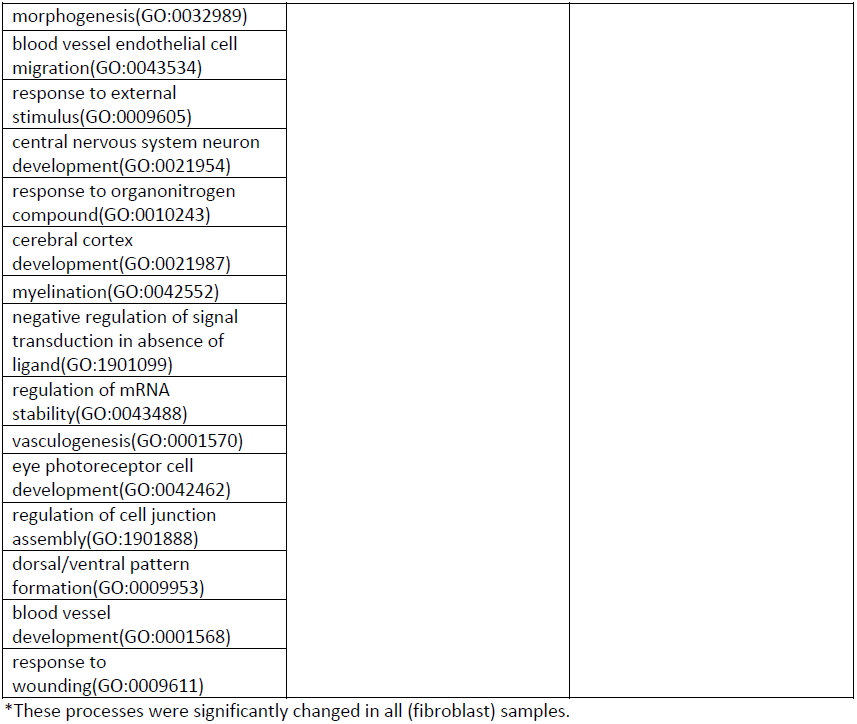
Changed biological processes for the different tissue groups, processes are found to be changed in at least two of three samples.

**Table 3:**
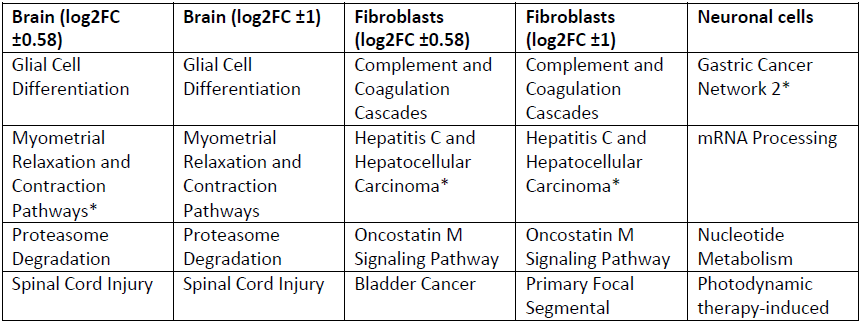
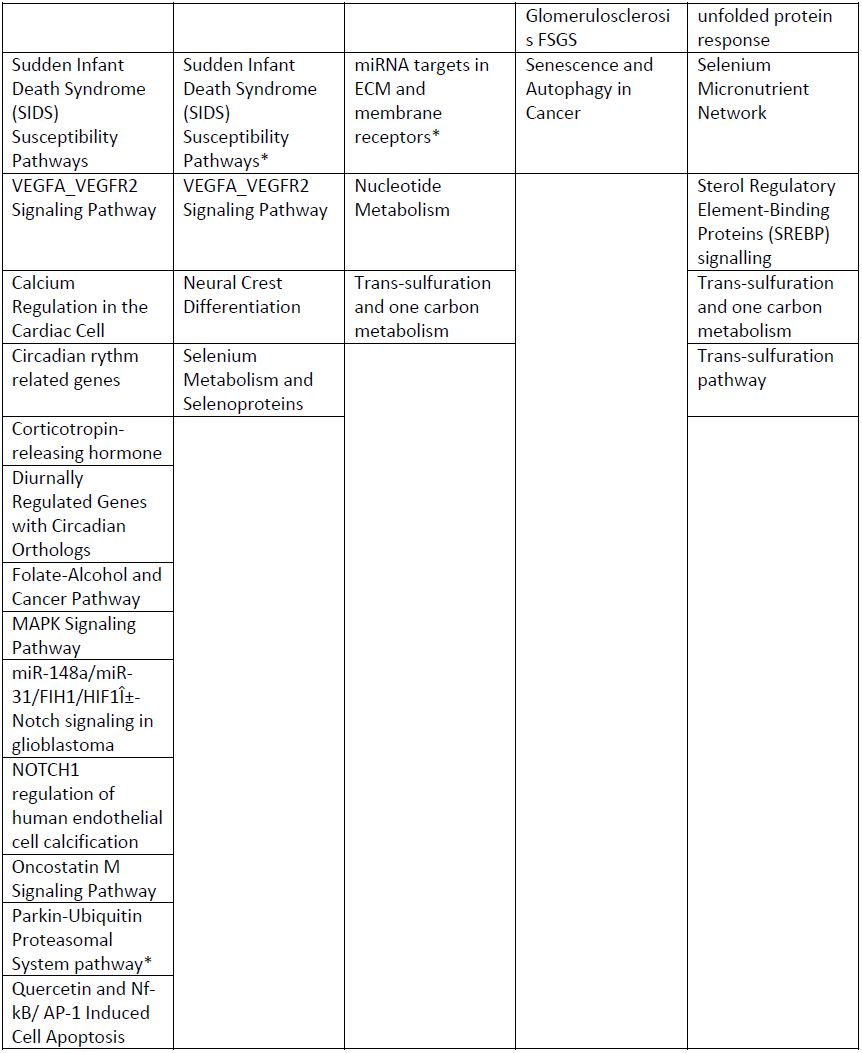
Molecular pathways commonly affected in the tissue groups

| **Brain (log2FC $\pm 0.58$)** | **Brain (log2FC $\pm 1$)** | **Fibroblasts (log2FC $\pm 0.58$)** | **Fibroblasts (log2FC $\pm 1$)** | **Neuronal cells** |
| --- | --- | --- | --- | --- |
| Glial Cell Differentiation | Glial Cell Differentiation | Complement and Coagulation Cascades | Complement and Coagulation Cascades | Gastric Cancer Network 2* |
| Myometrial Relaxation and Contraction Pathways* | Myometrial Relaxation and Contraction Pathways | Hepatitis C and Hepatocellular Carcinoma* | Hepatitis C and Hepatocellular Carcinoma* | mRNA Processing |
| Proteasome Degradation | Proteasome Degradation | Oncostatin M Signaling Pathway | Oncostatin M Signaling Pathway | Nucleotide Metabolism |
| Spinal Cord Injury | Spinal Cord Injury | Bladder Cancer | Primary Focal Segmental | Photodynamic therapy-induced |
|  |  |  | Glomerulosclerosis FSGS | unfolded protein response |
| Sudden Infant Death Syndrome (SIDS) Susceptibility Pathways | Sudden Infant Death Syndrome (SIDS) Susceptibility Pathways* | miRNA targets in ECM and membrane receptors* | Senescence and Autophagy in Cancer | Selenium Micronutrient Network |
| VEGFA_VEGFR2 Signaling Pathway | VEGFA_VEGFR2 Signaling Pathway | Nucleotide Metabolism |  | Sterol Regulatory Element-Binding Proteins (SREBP) signalling |
| Calcium Regulation in the Cardiac Cell | Neural Crest Differentiation | Trans-sulfuration and one carbon metabolism |  | Trans-sulfuration and one carbon metabolism |
| Circadian rythm related genes | Selenium Metabolism and Selenoproteins |  |  | Trans-sulfuration pathway |
| Corticotropin-releasing hormone |  |  |  |  |
| Diurnally Regulated Genes with Circadian Orthologs |  |  |  |  |
| Folate-Alcohol and Cancer Pathway |  |  |  |  |
| MAPK Signaling Pathway |  |  |  |  |
| miR-148a/miR-31/FIH1/HIF1±-Notch signaling in glioblastoma |  |  |  |  |
| NOTCH1 regulation of human endothelial cell calcification |  |  |  |  |
| Oncostatin M Signaling Pathway |  |  |  |  |
| Parkin-Ubiquitin Proteasomal System pathway* |  |  |  |  |
| Quercetin and Nf-kB/ AP-1 Induced Cell Apoptosis |  |  |  |  |

**Fig 1:**
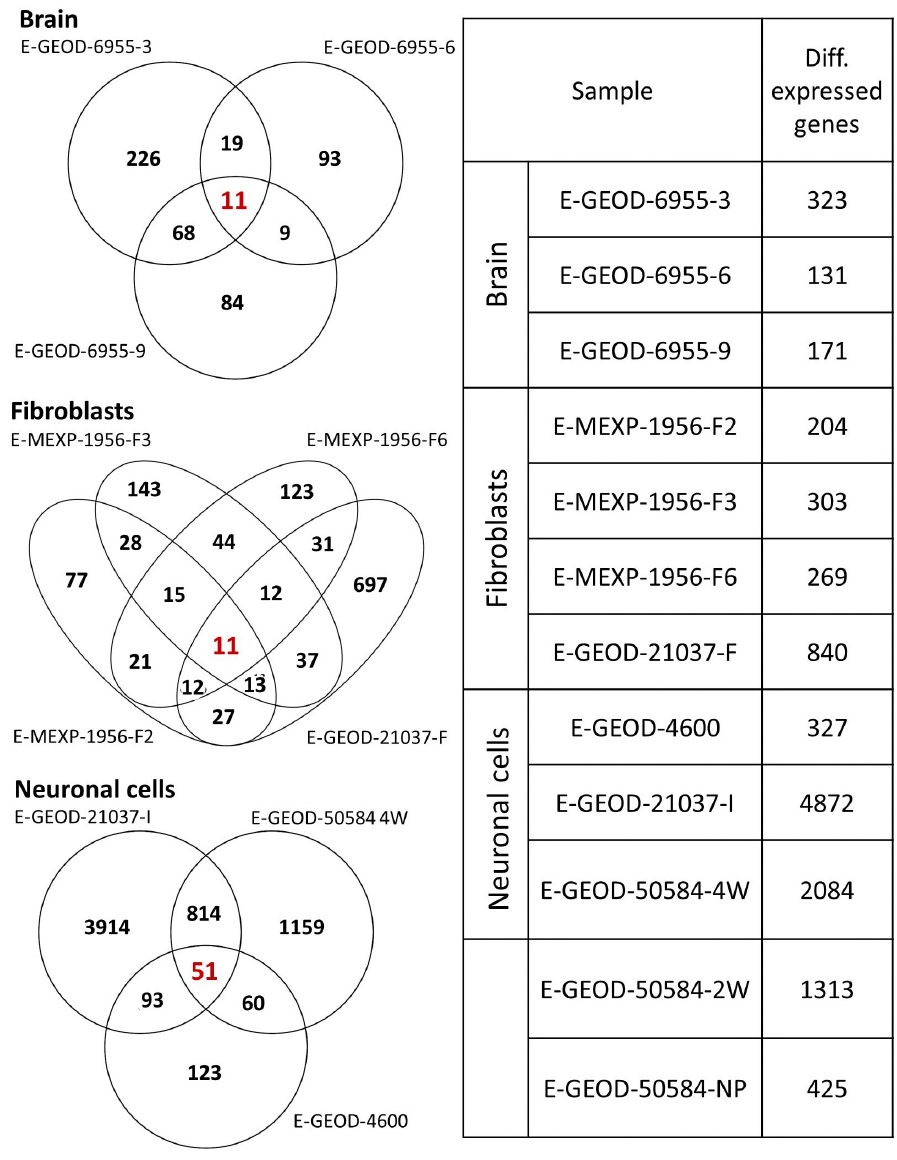
The overlapping differentially expressed genes in the three different tissue groups (red numbers) and the differentially expressed genes for each experimental tissue group (right). Log2FC threshold = ± 1

## GO analysis

The differentially expressed gene lists from each dataset were subjected to Gene Ontology (GO) analysis to identify enriched biological processes in which these genes are involved. These lists of changed GO processes (complete lists in sup. Table 4) were scanned for overlapping processes within the tissue group (Table 2). The results were “pruned” with GO-Elites algorithm to make sure subclasses with significant numbers of changed genes do not automatically lead to higher classes being found to be significantly changed, too.

**Table 4:** Selected transcriptomics datasets

| Dataset ID | Original study | Cell type used | Experimental group | MECP2 mutation | Array type |
| --- | --- | --- | --- | --- | --- |
| **E-GEOD-21037** | (Muotri et al, 2010) L1 retrotransposition in neurons is mediated by MeCP2 | Fibroblasts (untransfected) and induced pluripotent stem cells (iPSCs) from transfected fibroblasts | Fibroblasts<br>Cell culture (iPSCs)<br>Stem cells (iPSCs) | frameshift MeCP2 mutation | Affymetrix GeneChip Human Gene 1.0 ST Array |
| **E-MEXP-1956** | (Nectoux et al, 2010) Transcription profiling of cultured fibroblast cell lines from Rett syndrome patients | Human fibroblasts from skin biopsys, clones with either X(mutated) or X(normal) active. | Fibroblasts | Patient 1, p.R255X (nonsense), Patient 2, p.R270X (nonsense), Patient 3, c.1156del41 (frameshift) | Affymetrix GeneChip Human Genome U133 Plus 2.0 |
| **E-GEOD-6955** | (Deng et al, 2007) Transcription profiling of human super frontal gyrus (post mortem brain) from Rett syndrome patients | Human brain, super frontal gyrus samples from Rett patients and age matching control individuals | Brain | pool: 2 year old R168X + 4 year: n/a; 6 year: n/a; pool: 8 year R106W + 8 year R225X | Affymetrix GeneChip Human Genome U95Av2 |
| **E-GEOD-4600** | (Peddada et al, 2006) Transcription profiling by array of human SH-SY5Y cells transfected with an MeCP2 transcriptional repressor | "human SH-SY5Y neuronal cells transfected with a methylated oligonucleotide decoy to block MeCP2 binding to endogenous target genes during phorbol 12-myristate 13-acetate (PMA) induced maturational differentiation"(Peddada et al, 2006) | Cell culture | decoy to block MECP2 binding to targets | Affymetrix GeneChip Human Genome U133 Plus 2.0 |
| **E-GEOD-50584** | (Li et al, 2013)<br>Expression data from human ESCs-derived neural precursors (NPs) and neurons | WIBR cells - human ESCs, WIBR3 female, WIBR1 male, MECP2 mutation, precursor and after 2 and 4 weeks of differentiation | Cell culture<br>Stem cells | TALEN-mediated gene editing, targeting 3rd exon (MBD of MECP2 | GeneChip® PrimeView™ Human Gene Expression Array |

### Human brain

The major alterations in the brain affect several neurological processes such as regulation of synaptic transmission, substantia nigra development, neuron migration and behavior. Inflammatory and stress related processes are seen, too, e.g. hydrogen peroxide catabolic process, response to unfolded protein, response to wounding. Other processes are possibly due to the inclusion of blood and blood vessels like oxygen transport, blood vessel endothelial cell migration. DNA binding and mRNA stability are also affected processes.

### Fibroblasts

The fibroblasts of RTT patients show affected extracellular matrix organization and regulation of vascular endothelial growth factor production in all samples investigated. Other observed processes include response to stimuli (drug, dexamethasone, purine containing compound, organophosphorus, mechanical stimulus), and cell division.

### Neuronal cells

The neuronal cells derived from RTT patients, and the SH-SY5Y cell line, indicate significant affected processes within neuronal function (synapsis organization, spinal cord development, cerebral cortex development) but also in stress (response to UV), cell adhesion/cytoskeleton and blood vessel development. In the E-GEOD-4600 and in E-GEOD-50584 4W sample, but not significant in E-GEOD-20137, the translation process is strongly affected and especially translation initiation factors are dominantly downregulated (Fig 2).

**Fig 2:**
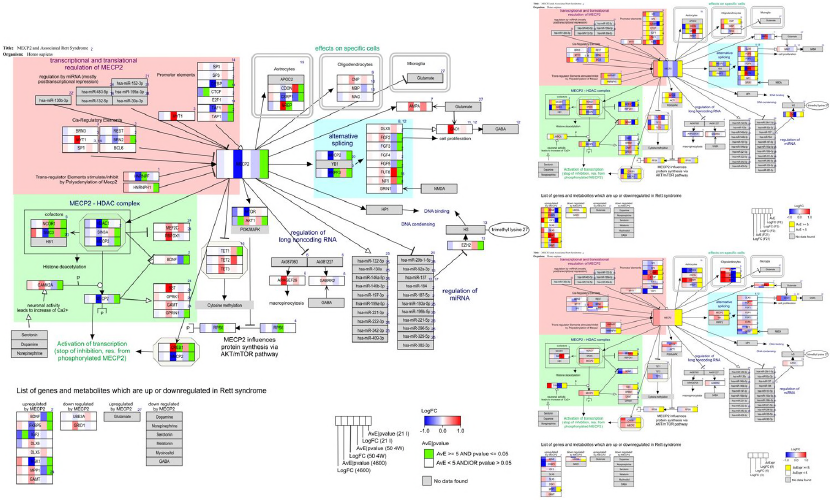
Significantly changed gene expression from E-GEOD 50584 4W (ESCs) and E-GEOD-4600 (SH-SY5Y) of the neuronal cell group which are related to mRNA processing and translation initiation factors. Note that, although the different expression level is lower in SH-SY5Y cells, in both datasets transcriptional initiation genes are dominantly down regulated in RTT samples.

## Pathway analysis

Pathway analysis was performed for each list of differentially expressed genes of the twelve datasets (and two different log2FC thresholds for the brain and fibroblast samples) to identify whether they affect certain biological pathways with known direct molecular interaction. Significantly changed pathways (thresholds: z-score ≥ 1.96 (equivalent to two standard deviations), p ≤ 0.05, minimum number of changed genes per pathway ≥ 3) were collected and analyzed (the complete list can be found in sup. Table 5, the overlapping pathways including z-scores in sup. Table 6 and the summary in sup. Table 3).

All samples showed energy and biosynthesis metabolism dysregulation in the RTT when compared to control samples: pathways affected were cholesterol/fatty acid metabolism, TCA cycle, cori cycle, amino acid metabolism, urea cycle, electron transport chain, or glycogen metabolism. Most samples show changes in at least one micronutrient involved pathways: zinc homeostasis, copper homeostasis, selenium, vitamin A, vitamin B12, folate, and iron metabolism. The most abundantly changed pathways (found in 5 of 12 comparisons - no pathway was found more often than in 5 samples (log2FC 1)) are: Gastric Cancer Network 2, Nucleotide Metabolism, Proteasome Degradation, Selenium Micronutrient Network, Trans-sulfuration and One carbon metabolism. Pathways changed in 4 of 12 samples are: Apoptosis-related network due to altered Notch3 in ovarian cancer, Gastric Cancer Network 1, Hepatitis C and Hepatocellular Carcinoma, mRNA Processing, Oncostatin M Signaling Pathway, Parkin-Ubiquitin Proteasomal System pathway, Photodynamic therapy-induced unfolded protein response, Retinoblastoma (RB) in Cancer, Selenium Metabolism and Selenoproteins, Spinal Cord Injury, and VEGFA-VEGFR2 Signaling Pathway.

Using the threshold of 0.58 for log2FC revealed more affected pathways (as there were also more by definition differentially expressed genes) nevertheless, the tendency was unaffected. The pathways for each single dataset were then investigated for overlapping pathways between the datasets of one tissue group (Table 3).

Brain tissue: In brain tissue, the affected pathways were mainly involved in neurological processes like Glial Cell Differentiation, Spinal Cord Injury, SIDS susceptibility but also in Proteasome Degradation and endothelial development (VEGFA-VEGFR2 Signaling pathway). For the lower threshold of 0.58 also Circadian rhythm related genes, apoptosis and inflammation related pathways show up while within the gene list derived with the more restrictive threshold another neuronal development pathway (Neuronal Crest Differentiation) and the Selenium Metabolism and Selenoproteins pathway showed up.

Fibroblasts: In the fibroblast samples, there were dominantly cell cycle related pathways found to overlap between the single datasets. The lower threshold dataset also revealed changes in Extracellular Matrix (miRNA targets in ECM and membrane receptors), Nucleotide and Trans-sulfuration and one carbon metabolism.

Neuronal cells: Although the single datasets had a lot of changed pathways in neuronal and neurodevelopmental processes, the overlapping pathways between the three datasets were involved in cell cycle (Gastric Cancer Network 2), mRNA processing and translation factors, nucleotide metabolism, stress and apoptosis (Photodynamic therapy-induced unfolded protein response), SREBP signaling, and trans-sulfuration pathways.

### Pathway visualization

To visualize the direct molecular environment of MECP2 the pathway “MECP2 and associated Rett syndrome” was used as described previously (Ehrhart et al, 2016). *BDNF, FKBP5* and *DLX6* were found to be downregulated in several but not in all samples while *GRID1* was upregulated but found only in one single sample (E-GEOD-50584 4W). For *DLX5, SGK1* and *UBE3A* there was no significantly changed gene expression found in any of the samples. *GAMT, IGF2, MPP1* and *FXYD1* were upregulated although expected to be downregulated ((Ehrhart et al, 2016) and literature cited there). *GAMT* was found to be upregulated with log2FC of about 0.3 only in the ESC (E-GEOD-50584) samples. *IGF2* was downregulated in human fibroblasts (E-MEXP-1956-F6) and SH-SY5S cells (E-GEOD-4600) while being strongly (log2FC = 6) upregulated in another sample of human fibroblasts (E-GEOD-21307-F). *MPP1* was upregulated in two of three human fibroblast samples (both log2FC > 1) but slightly downregulated in iPSCs (E-GEOD-21307-I) (log2FC = -0.57). *FXYD1* was found only in human brain samples and upregulated with log2FC 1.20 – 1.52, but as the average expression was below the threshold of 5 these were not counted as relevant.

*MECP2* transcript itself is significantly lower expressed in all RTT neuronal cell culture samples (including the precursor samples E-GEOD-50584 NP and 2W) and three of four fibroblast samples (all MEXP-1956 samples) with a log2FC between -0.1 (E-GEOD-4600) to -1.5 (E-GEOD-50584 2W). In human brain samples the *MECP2* transcript levels are slightly increased (log2FC 0.2 - 0.4) as well as in E-GEOD-21037 fibroblasts (0.4). For the reliability of the samples, the cell culture samples provided biological replica (it was possible to calculate a p-value, although this is of limited added value with only 2 replica) and cell culture samples had the advantage that their gene expression is more similar than comparing different individuals. This is also seen in some of the fibroblast samples. For the study E-MEXP-1956, each of the three RTT and control pairs were derived from the same individual but clones with different X chromosomes active.

MECP2 is supposed to inhibit expression of MEF2C and RBFOX1 via DNA compaction (MECP2-HDAC complex). The dysfunctional *MECP2* expression profile shows a slight but in cell culture samples significant upregulation of these genes (Fig 3). Those genes, which are supposed to be only activated when MECP2 is phosphorylated and therefore detached from the DNA to allow RNA transcription, are here observed to be higher expressed: *SST, OPRK1, GAMT*, and *GPRIN1*. This indicates that MECP2 does not block their transcription anymore. Influence on AKT/mTOR pathway and TET complex seems indifferent. In the fibroblast cell samples there is no visible tendency of an effect in the MECP2 regulated genes (Fig 3, lower right).

**Fig 3:**
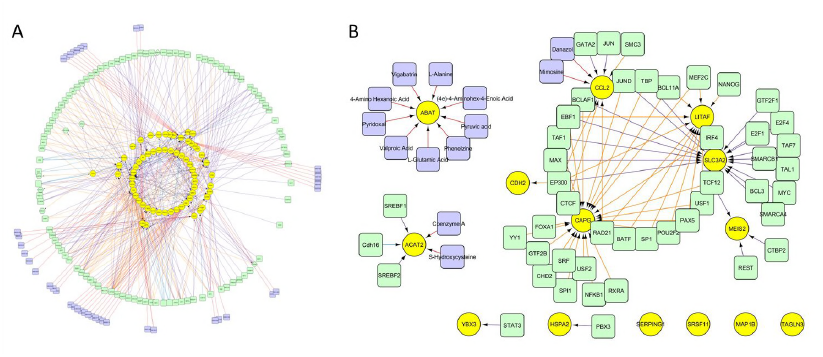
MECP2 pathway including data from the different tissue groups. **Neuronal cells:** Each gene box indicates log2FC (blue – red) and significance (if p < 0.05). **Brain:** Each box contain log2FC (from -1 (blue) to 1 (red)) for the 3 different age groups and average expression level (yellow if > 5). Horizontal separation in a box indicates multiple Affymetrix identifiers (probes) which are linked to the same gene. **Fibroblast:** Each box contain log2FC (from -1 (blue) to 1 (red)) for the 4 different fibroblast samples and their average expression level (yellow if > 5). High resolution images of the single pathways are in the supplementary data.

## Network extension

Network analysis approaches, using Cytoscape, were used to visualize the genes of interest and extend the results of the GO analysis by integration of external database knowledge.

### Analysis of the genes of interest

In a first step, the list of genes, which were identified as consistently differentially expressed one tissue group and in at least one more dataset (the 14 genes from Table 1 and the 121 from sup. Table 2 for a different threshold) were imported to Cytoscape as data nodes. Then the network was extended using CyTargetLinker app for Cytoscape and the *Homo sapiens* package of RegINs containing transcription factors from ENCODE and drug targets from DrugBank.

The result of network extension is shown in figure 4. Several transcription factors are already known to interact with MECP2, e.g. E2F1, MEF2C, REST, SIN3A, SMC3, SP1, and TAF1 (full list of added transcription factors and drugs in supp. Table 7). Recognized as drug-targets were ACAT2 (S-Hydroxycysteine and Coenzyme A, which are rather substrates than drugs), CCL2 (Mimosine and Danazol, both inhibitors and approved drugs), and ABAT (Valproic Acid, Vigabatrin, and Phenelzine, these are approved drugs and inhibitors to the target; Pyridoxal Phosphate and Pyruvic acid, - nutraceuticals, inhibitors; (4e)-4-Aminohex-4-Enoic Acid and 4-Amino Hexanoic Acid – these are experimental drugs; L-Glutamic Acid, and L-Alanine).

**Fig 4:**
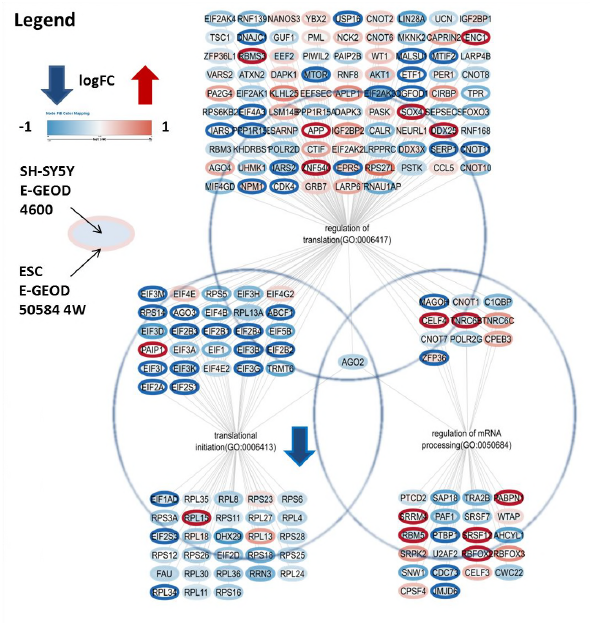
Network extension for the differentially expressed genes (yellow) using data from ENCODE proximal and distant transcription factors (light green), drug targets and drugs (violet) for (A) threshold absolute log2FC > 0.58 identified gene list and (B) absolute logFC > 1 gene list.

## Discussion

The aim of the study was to establish a workflow to analyze multiple raw transcriptomics datasets with a common experimental setup (namely, human sample with impaired *MECP2* vs. control) and to investigate the differentially expressed genes for RTT on the level of genes, biological pathways and processes. The approach was a data driven identification of changed gene expression profiles and inclusion of prior knowledge (WikiPathways, GO, ENCODE, DrugBank) for the analysis of the biomedical impact in RTT. For an integrative study, we are dependent on experimental execution and data quality of the original studies, and the correct and extensive annotation of data. Having one microarray per sample – without technical and biological replica – is hardly enough to draw conclusions, however data integration with similar studies, can help to overcome the problem of too few samples if the biological and experimental backgrounds are similar.

### Gene expression with impaired MECP2 – hypothesis vs. data

Genes, which are involved in RTT disorder development and progression, according to previously published studies are *MECP2, CDKL5, EGR2, PTPN1, BDNF, GAMT, FKBP5, IGF2, DLX5, DLX6, SGK1, MPP1, FXYD1, UBE3A*, and *GRID1* (Ehrhart et al, 2016). These genes were found again in the data but the reported outcome shows a greater variability than reported in literature. *BDNF* e.g. is mostly reported to be down regulated in RTT (or model systems). We found a significant downregulation only in one fibroblast and one iPSC sample, in other samples gene expression changes were not significant and occurred in both directions. We did not find a significant change in expression profile for either *DLX5* or *DLX6* and can therefore support the hypothesis that it is not involved in RTT disorder progression (Miyano et al, 2008; Schule et al, 2007). A general observation was that the gene expression patterns vary greatly among the different samples which matches previous observations e.g. from (Tanaka et al, 2014). This study investigated iPSCs derived from RTT patients (and a clone with healthy X-chromosome as control) and all of them had different causative *MECP2* mutations. The result was that gene expression profiles for those different patients were very different. In our study, too, there is no single mutation occurring twice in the sample datasets. Low overlap of previously known genes and identified by the data-driven approach was reported several times before (e.g. epilepsy study (Rogic & Pavlidis, 2009)) and all these studies revealed new genes which were not known to be involved in disease progression, yet (Ch’ng et al, 2015; Mistry et al, 2013; Raddatz et al, 2014; Roder et al, 2012).

The choice of thresholds is a critical issue in this study. A significance threshold in form of a p-value could only be used if a certain amount of biological replicas were available. If there were only one sample or only technical replicas available the threshold was defined by log2FC of either > 1 or <-1 (res. 0.58) - meaning double or 1.5fold expression level than control. Several smaller but important changes in gene expression therefore may not be detected. We evaluated a higher and a lower threshold for those samples where no p-value could be calculated. Although, the higher threshold resulted in a smaller list of genes, the list of pathways and processed they were involved remained similar. Also the platform specific average expression level was used as threshold between relevant and not relevant. This is a criterion which is good for artificial systems like the cell cultures but possibly for human derived tissue too stringent. *FXYD1* e.g. was indeed differently changed with log2FC between 0.7 and 1.51 but due to low average expression we did not count it as differentially expressed whereas the original authors did (Deng et al, 2007).

### Newly identified differentially expressed genes are relevant in known RTT pathways

After grouping the available experimental data in three sub-groups (brain tissue, fibroblasts, and neuronal cells) we identified several genes, which were not yet mentioned before in relation to RTT. Analyzing their function in the context of pathways (molecular interaction) and function (functional connection via GO) shows that they are clearly involved in functions and pathways known to be involved in RTT disease progression and therefore deserve further attention.

The changes in the expression of genes from human brain samples were mostly related to regulation and signal transduction pathways as well as response to chemical stress which may be due to apoptosis related processes. *LITAF* (which is found to be significantly differentially expressed in eight independent datasets for threshold log2FC 0.58 and in six for log2FC 1) is lipopolysaccharide induced and *HSPA2* usually is stress-induced. Inflammation (and energy metabolism) pathways were also found in the other tissue groups and described for RTT before (Pecorelli et al, 2016), so inflammation related processes may be involved in progression of the disorder itself. The human brain dataset is the most interesting tissue for the study of RTT, but needs to be interpreted with care. Human brain tissue is usually only available postmortem and the process of dying is mirrored in the gene expression profiles. Also the sample extraction concerning the timing and cooling chain is a potential cause of problems especially for easily degradable RNA samples. For these samples the extraction was performed between 2.9 and 36 h postmortem (supplementary material of (Deng et al, 2007)) and our quality control showed no noteworthy RNA degradation. Another cause of detection of inflammatory pathway activities in brain tissue (or primary tissue in general) might be the extraction of RNA from white blood cells present in the tissue.

Human brain samples showed no affected pathways or processes concerning energy metabolism pathways, whereas all other samples do. This is possibly a cell culture effect, that growing and actively differentiating cells may have a higher energy need than mature neurons in a human brain and therefore are able to indicate the differences between MECP2-deficient and control status.

Human brain and fibroblasts revealed the greatest changes in micronutrient related pathways, namely calcium, copper, selenium, vitamin A and zinc (brain) and copper, folate, iron, and Vitamin B12 (fibroblasts). As mal- and undernutrition is often observed in RTT females and since both brain and fibroblast samples are the ones which are most directly taken from human patients this may reflect their nutritional problems. These changed pathways do not show up so much in cell culture samples which might be due to the culture conditions, namely full nutrient supply by complete growth medium. Investigation where in the process between eating, digestion, uptake in blood and cellular delivery the problems occur in RTT females might help to understand some of the symptoms. This analysis may also indicate that the cells are able to take up the nutrients and use them once they are delivered so the error must happen before. Further studies of gut and nutrient processing tissues of RTT females and model systems may be helpful to elucidate this issue.

Changes in cell cycle related pathways are found in all samples but mostly in fibroblasts and least in human brain. This may again be a result of the higher sensitivity of cell culture systems because of the higher metabolic activity in cell culture procedure and influence of growth factors in cell culture medium but still indicates that the influence of MECP2 reaches to the regulation of cell cycle in different degrees in the different samples.

The typical RTT or MECP2 related pathways are not significantly changed in fibroblasts. This is expected as these neuronal pathways are not active in fibroblasts. But still, there are changed developmental pathways and processes in fibroblasts as well, e.g. cell differentiation process. *MECP2* seems to be involved in cellular function not only in neuronal cells. This matches the data given on tissue atlas that *MECP2* RNA and protein are found in almost any tissue with yet unknown function. However, one of these functions in fibroblasts seems to be related to the cell cytoskeleton, adhesion and extracellular matrix, which are most affected in fibroblasts indicating that *MECP2* is interfering with this process also in non-neuronal cells as well. Cellular compartment organization, e.g. actin/microtubule and cell adhesion processes, may link to *MECP2* indirectly influencing the directed outgrowth of neurites in brain tissue samples and neuronal cell culture and of filopodia of fibroblasts. Differentiation related pathways show the biggest differences in human brain and the least in fibroblasts and suits the hypothesis that human RTT brain is immature. Cell culture samples of cells which are on a differentiation protocol towards neuronal cells show an intermediate state between fibroblasts and human brain. Microtubules are deacetylated too much in RTT due to HDAC6 dysregulation which weakens the cytoskeleton. Addition of Tubastatin A ameliorates the effect in cell culture of fibroblasts and SH-SY5Y cells (Gold et al, 2015). Changes in the ratio of deposition of collagen I and III was observed but this seems to be a subclinical issue of the disorder as no skin, wound healing problems, or premature aging are reported yet in RTT females (Signorini et al, 2014).

### Translation and RNA processing is impaired in RTT

Changes in RNA processing are identified clearer by GO analysis than pathway analysis because in the pathways from WikiPathways database the changed genes are either “hidden” in non-significant pathways or not yet included (WikiPathways currently includes about 7000 human genes). MECP2 is a known regulator of transcription and translation both generically and for single genes and the data shows that mRNA processing is altered in RTT and especially genes involved in translation initiation are massively downregulated (Fig. 5). *MTOR* is also downregulated which fits the observation that the total protein levels of RTT cells is lower (Ricciardi et al, 2011). The AKT/mTOR pathway is a known disease pathway for neurofibromatosis (Banerjee et al, 2011).

**Fig 5:**
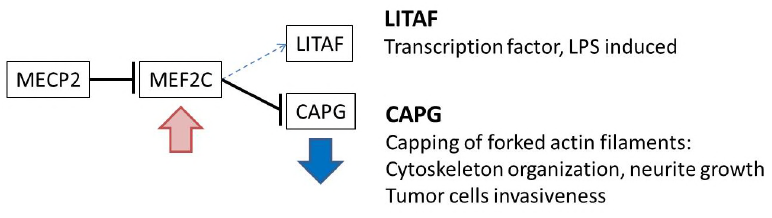
MECP2 wildtype inhibits expression of MEF2C which inhibits expression of CAPG. If MECP2 is dysfunctional, CAPG was found to be mainly downregulated in the expression data. As MECP2 inhibition of MEF2C does not work anymore, MEF2C level increases (insignificant trend to upregulation found in data) and CAPG expression is inhibited. MEF2C is also a transcription factor for LITAF but the effect (stimulation or inhibition) is not clear from the data.

Alternative splicing is also changed in RTT, there are several genes which are known to be differentially spliced in RTT (or MECP2 deficient models), namely *DLX5* (a transcription factor which stimulates expression of *GAD1* (GABA receptor)) (Miyano et al, 2008), *FGF2* (a cell proliferation factor), *FGF3, FGF4, FGF5, FUT8, NF1* (neurofibromatosis 1), and *GRIN1* (NMDA receptor) (Cartron et al, 2013; Long et al, 2011; Nomura et al, 2008; Smith & Sadee, 2011; Young et al, 2005). Differential gene expression of splicing factors was also identified in a transcriptomics meta-study in schizophrenia (Mistry et al, 2013). There are multiple splicing factors differently expressed which may be the cause of various splice variants observed in RTT. The spliceosome includes SR proteins (differentially expressed in this study: *SRSF2, SRSF5, SRSF7*) which are adaptors which link mRNA to NXF1 (nuclear export factor 1) for alternative RNA processing and export (Muller-McNicoll et al, 2016) and help to form the spliceosomal complex A (WP411). SRPK2 is a SR protein kinase. PRPF8, SF3B5, SNRPB, SNRPB2 SNRPD2, SNRPF, are essential parts of the spliceosome components. CLP1 is a component of tRNA endonuclease complex (TSEN) complex and an essential part for tRNA maturation. Mutations in *CLP1* have been proven to cause neurodegenerative disorders (Schaffer et al, 2014). RBM5 belongs to the family of RNA-binding proteins, which are known as splicing regulators. PTBP1 and PTBP2 bind pre-mRNA and direct the spliceosome attachment process to the correct site. SUGP2 is a (mRNA binding) splicing factor. SF3B1 is a subunit of splicing factor SF3B. PABPN1 is a poly(A) binding protein and involved in mRNA3’ end processing. Differentially expressed genes responsible for “RNA splicing(GO:0008380)” in the group neuronal cells are *PRPF38B, SNRPB2*, and *SRSF11*.

Bedogni and coworkers (Bedogni et al, 2014) had a look at animal studies in RTT and found that (apart from several cytoskeleton related and cell metabolism related pathways) intracellular signaling, namely the eukaryotic initiation of transcription factors (EIF) signaling pathway, was affected. Many EIFs which are found in this study to be differentially expressed, respectively, downregulated. Impaired function of EIF pathways regulating protein production was associated with several disorders (Chang et al, 2006). EIF2 was previously identified as one of the factors involved in age-associated spatial learning impairment – also identified by a transcriptomics meta-study (Uddin & Singh, 2013). Therefore our data supports the hypothesis that EIF signaling and regulation pathways are relevant causing the disorder phenotype in RTT, too.

### Possible connection between MECP2 and CAPG via MEF2C

Using network extension we investigated whether there are known transcription factors which may explain the regulation of the genes and processes back to their core gene MECP2. Several transcription factors (proximal and distal) which were added by network extension are already known to interact with MECP2: E2F1, MEF2C, REST, SIN3A, SMC3, SP1, and TAF1. E2F1 and TAF1 are promotor elements of MECP2 ((Ehrhart et al, 2016) and literature cited there). SP1 is known as cis regulatory element and promotor element of MECP2. REST is a cis regulatory element of MECP2. SIN3A is an integral part of the MECP2-HDAC complex and SMC3 is one of the cofactors of this complex. *MEF2C* expression is inhibited by (functional) MECP2 together with HDAC complex. It interacts with EP300, several histone deacetylases, and SP1. Mutations of MEF2C are known to cause mental retardation and psychomotor impairment (Le Meur et al, 2010). Furthermore, the network extension with known transcription factors revealed a possible connection between cytoskeleton organization and MECP2 via MEF2C and CAPG. We hypothesize that MECP2 regulates CAPG expression via MEF2C (Fig 5).

These results, nevertheless, need to be handled with caution because the number of studies and samples often is limited and the setting of thresholds is crucial. More data, integration of data (e.g. from animal studies or other kinds of omics data) will allow to refine the results. Our analysis revealed genes which were yet unknown to be involved in disease progression but often related to processes which are known to be affected. They are also often found to be affected in other mental disorders. A general conclusion of our analysis is that “one gene – one disease” is definitively true as a basis for RTT, but the link between mutation and phenotypic outcome is defined by more than this one gene. This connection involves a complex network in which individual genetic variation and environmental influence (e.g. nutrition) determine the disorder outcome.

## Materials and Methods

### Transcriptomics datasets

All publically available MECP2 related transcriptomics datasets were chosen from open access databases, namely GEO (Edgar et al, 2002) and ArrayExpress (Rustici et al, 2013), and used in the present study: E-GEOD-21037, E-MEXP-1956, E-GEOD-6955, E-GEOD-4600, and E-GEOD-50584. The study selection protocol (including PRISMA flow diagram (Moher et al, 2009)) and details about the datasets can be found in supplementary data (Supp. Table 1 Dataset description, selection criteria and Sup Figure PRISMA flow diagram). The study details and publications are listed in Table 4. In total five different studies were selected from which 48 samples were included in the present study. In total, in the five studies 24 microarrays with an impaired *MECP2* gene and 24 matching controls were found.

The protocol for the formation of the experimental groups and the processing of each study was as follows:

#### E-GEOD-21037 fibroblasts and fibroblast derived iPSCs (Muotri et al, 2010)

RTT and WT fibroblasts were obtained from two different individuals - one with Rett syndrome and one healthy person. For untransfected fibroblasts there were three technical replicas. The mean value of the replica was used for log2FC calculation. For iPSCs there were three technical replicas of two different clones given. The clones were treated as biological replica. The mean value of each clone was used for log2FC and p-value calculation using arrayanalysis.org statistics module. HUES6 is a control stem cell line and the data was not used for this study.

**➔ Experimental groups: E-GEOD-21037-F and E-GEOD-21037-I**

#### E-MEXP-1956 human primary fibroblasts (Nectoux et al, 2010)

The fibroblast samples were taken from three individuals: F2, 3 and 6. For each individual RTT vs. WT was calculated separately and log2FC > |1| and log2FC > |0.58| was used as threshold. The choice of RTT and WT samples was performed according to the study authors: ‘Transcriptomic analysis was indeed performed with six clones: one matched pair (one mutant and one wild-type clone) from each of the three *MECP2* mutation patients. When several wild-type or mutant clones were available from the same patient, the one presenting the closest 100:0 skewed X-inactivation score was selected for the transcriptomic analysis.’

**➔ Experimental groups: E-MEXP-1956-F2, F3 and F6**

#### E-GEOD-6955 human brain samples (Deng et al, 2007)

Human postmortem brain samples from the superior frontal gyrus were used for this study. The samples were originally derived from a human tissue bank and neither the cause of death nor the circumstances of sample collection were available for this study. There was only one sample per age group (3, 6, and 9 years) and two of the RTT samples are pooled from two patients. We compared the age groups separately and used log2FC > |1| and log2FC > |0.58| as threshold. The original dataset provided Affymetrix identifiers. These were converted to Ensembl identifiers using Biomart.

**➔ Experimental groups: E-GEOD-6955-3, E-GEOD-6955-6 and E-GEOD-6955-9**

#### E-GEOD-4600 SH-SY5Y neuroblastoma cell line (Peddada et al, 2006)

SH-SY5Y neuroblastoma cells were transfected with either a MECP2 decoy or a control decoy and differentiated towards neurons. The three samples of each group are biological replica. We used arrayanalysis.org statistics module to calculate log2FC and p-value. The samples from undifferentiated and differentiated wildtype SH-SY5Y cells were not used in this study. We considered the control decoy the most suitable control for this study.

**➔ Experimental group: E-GEOD-4600**

#### E-GEOD-50584 human ESCs (Li et al, 2013)

The authors of the original study gave no explicit information whether the samples are biological or technical replica. Due to sample naming and being cell cultures we assumed biological replica and calculated log2FC and p values using arrayanalysis.org statistics tool. We compared the three differentiation status (neuronal precursor (NP), after two (2W) and four weeks (4W) of differentiation) separately and used only the female cell samples (WIBR3). P value < 0.05 was used as threshold.

**➔ Experimental groups: E-GEOD-50584-NP, E-GEOD-50584-2W and E-GEOD-50584-4W**

### Analysis workflow

An analysis workflow was designed to compare and integrate the experimental groups on the level of genes and pathways (Fig 6). First, the changed gene expression profiles or pathway profiles were extracted. Second, these profiles were grouped and analyzed according to tissue or sample similarity into three main groups: human brain tissue, fibroblasts, and neuronal cells.

**Fig 6:**
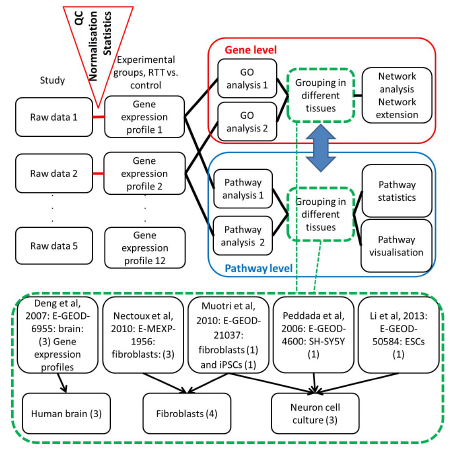
Workflow to integrate and analyze the MECP2-related transcriptomics datasets. Analysis of raw data results in a list of significantly (above the threshold) changed genes. Datasets were grouped according to their tissue origin into Human brain, Fibroblasts, and Neuron cell culture. Changed gene lists are the basis for a gene-wise comparison or search for commonly differentially expressed genes compared directly (using GO analysis) or a pathway approach where gene lists are first used to identify affected molecular pathways and then the commonly differentially affected pathways are identified.

### Quality control and data normalization

Raw transcriptomics datasets (.CEL files) were downloaded from GEO (Edgar & Barrett, 2006) or ArrayExpress (Kolesnikov et al, 2015), respectively. Quality control and GC-RMA normalization were performed using the ArrayAnalysis.org web tool (Eijssen et al, 2013). All datasets selected passed the quality control. The detailed QC reports can be found in supplementary data. For annotation of data Ensembl gene identifiers were used.

### Statistics

The ArrayAnalysis.org statistics module was used to perform statistical calculations (an adjusted t-test using limma package of Bioconductor/R) on the normalized data of those datasets where more than one biological replica was available (namely E-GEOD-21307-I, E-GEOD-4600, and E-GEOD-50584 all groups, see also details below). Fold change (FC), log2FC, average expression and p-values were calculated comparing RTT/MECP2 deficient-sample versus control. For those samples with only one (biological) replica we calculated log2FC and average expression (Rett/MECP2 deficient – WT/control): E-GEOD-21307 fibroblasts, E-MEXP-1956 fibroblasts, and E-GEOD-6955 human brain. Genes were considered as differentially expressed if the average expression is higher than 5 (which is recommended for the Affymetrix micro array platforms used), and if the p-value is smaller than 0.05. If the p-value was not available (for the human brain and fibroblast samples) log2FC > |1| or log2FC > |0.58| was taken as criterion.

To compare the various lists of differentially expressed genes the online tool Venny (version 2.1) 35 was used. We defined four different tissue groups (see Fig 6) i) Human brain: E-GEOD-6955: three different age groups of 3, 6 and 9 years, ii) Fibroblasts: E-GEOD-21307-F, E-MEXP-1956-F2, F3 and F6, iii) Neuronal cell culture (stem cells, which are differentiated towards neurons): E-GEOD-4600, E-GEOD-21307-I and E-GEOD-50584-4W, iv). Per tissue group the overlapping differentially expressed genes were extracted.

### Gene ontology analysis

Gene ontology (GO) analysis (overrepresentation analysis, ORA) was performed using GO-Elite (Version 1.2 Beta) (Zambon et al, 2012). Biological process was the GO category used in this analysis. The number of permutations for ORA was set to 2000. A z-score ≥ 1.96, permuted p-value ≤ 0.05 and minimum number of changed genes of 3 were set as cut-off and the results were pruned. For analysis the list of differentially expressed genes per sample dataset was used (criteria see above). As background, the lists of all genes measured were merged. Duplicates and identifiers, which were not recognized, were removed from the lists.

### Pathway analysis

PathVisio (version 3.2.0) (Kutmon et al, 2015) was used to visualize log2FC and p-values and to perform an overrepresentation analysis with the pathway statistics module using the lists of differentially expressed genes. For identifier mapping BridgeDb (version 82) (van Iersel et al, 2010) was used and as pathway database WikiPathways, namely the *Homo sapiens* curated pathway database, downloaded 17.08.2016, was used. The list of differentially expressed genes per sample dataset was used as input (criteria see above). A pathway was identified as changed if z-score ≥ 1.96, permuted p-value ≤ 0.05 and the number of genes in the pathway which meet the criteria ≥ 3.

### Network visualization and extension

Based on the obtained GO terms and gene lists, networks were generated using the network analysis tool Cytoscape (version 3.4.0) (Shannon et al, 2003; Smoot et al, 2011). The Cytoscape app, CyTargetLinker (Version 3.0.1) (Kutmon et al, 2013) (including RegIN file set for *Homo sapiens*) was used to extend the previously described network and add database information. The RegINs used are derived from ENCODE (transcription factors, proximal and distal) and DrugBank (drug targets).

## Acknowledgements

The authors would like to thank the authors of the original studies for providing their data in a FAIR way, making these kinds of meta-studies possible. FE and NS are funded from grants of Stichting Terre (Rett Syndroom Funds).

## Authors contributions

FE – study design, study execution, paper writing, SC – advice in analyzing the original studies and design of experimental groups, critical discussion, EC – pathway ontology analysis, critical review, LE – advice in statistics and design of experimental groups, ES – advice in clinical aspects of RTT, critical review, NS – critical review, CE – study design, critical discussion and review, LC – advice in study design, critical discussion and review

